# Safe plant Hsp90 adjuvants elicit an effective immune response against SARS-CoV2 derived RBD antigen

**DOI:** 10.1101/2023.11.25.568642

**Authors:** Victor A. Ramos-Duarte, Alejandro Orlowski, Carolina Jaquenod de Giusti, Mariana G. Corigliano, Ariel Legarralde, Luisa F. Mendoza-Morales, Agustín Atela, Manuel A. Sánchez, Valeria A. Sander, Sergio O. Angel, Marina Clemente

## Abstract

To better understand the role of pHsp90 adjuvant in immune response modulation, we proposed the use of the Receptor Binding Domain (RBD) of the Spike protein of SARS-CoV2, the principal candidate in the design of subunit vaccines. We evaluated the humoral and cellular immune responses against RBD through the strategy “protein mixture” (Adjuvant + Antigen). The rRBD adjuvanted with rAtHsp81.2 group showed a higher increase of the anti-rRBD IgG1, while the rRBD adjuvanted with rNbHsp90.3 group showed a significant increase of anti-rRBD IgG2b/2a. These results were consistent with the cellular immune response analysis. Spleen cell cultures from rRBD+rNbHsp90.3-immunized mice showed significantly increased IFN-γ production. In contrast, spleen cell cultures from rRBD+rAtHsp81.2-immunized mice showed significant increased IL-4 levels. Finally, vaccines adjuvanted with rNbHsp90.3 induced higher neutralizing antibody responses compared to those adjuvanted with rAtHsp81.2. To know whether both chaperones must form complexes to generate an effective immune response, we performed co-immunoprecipitation (co-IP) assays. The results indicated that the greater neutralizing capacity observed in the rRBD adjuvanted with rNbHsp90.3 group would be given by the rRBD-rNbHsp90.3 interaction rather than by the quality of the immune response triggered by the adjuvants. These results, together with our previous results, provide a comparative benchmark of these two novel and safe vaccine adjuvants for their capacity to stimulate immunity to a subunit vaccine, demonstrating the capacity of adjuvanted SARS-CoV2 subunit vaccines. Furthermore, these results revealed differences in the ability to modulate the immune response between these two pHsp90s, highlighting the importance of adjuvant selection for future rational vaccine and adjuvant design.

## 1. Introduction

During the pandemic of COVID-19 that occurred in the 2019-2022 period, researchers demonstrated that vaccination is one of the most successful strategies to prevent infectious diseases expansion in the human population, including those caused by emerging viruses[1]. At record-breaking speed, the scientific community has worked jointly to develop different vaccine formulations that contribute to the protection of humanity[2]. However, some needs persist to improve existing vaccines or produce new ones that guarantee the vaccination of the entire world population with democratic and broad access, especially in emerging countries. Likewise, the appearance of new variants continues today, and the rational design of vaccines that adapt to these new COVID-19 variants or those that may arise in the future will continue to be essential to achieve greater efficacy and a long-lasting period of protection[3]. Therefore, it is clear that the future challenge is to design a wide variety of alternatives to anti-COVID-19 vaccines or vaccination strategies that guarantee high tolerability and low side effects while avoiding reactogenicity. In this regard, subunit vaccines using proteins or peptides are much safer as they do not contain any live viral components and do not cause severe undesirable side effects[4]. However, subunit vaccines derived from known pathogen target antigens are generally less immunogenic, which can be improved using appropriate adjuvant(s) to the vaccine[1].

Adjuvants not only enhance the immunogenic capacity of the vaccine formulation, but also contribute to decreasing the amount of antigen needed for each vaccine dose or reducing the number of vaccines doses[5]. In addition, adjuvants stimulate a Th2 biased immune response, enhancing antibody production[6]. In this sense, the Hsp90s have been extensively studied, and their immunomodulatory properties have contributed significantly to improving vaccine development against infectious diseases, especially against intracellular pathogens[7]. The HSP90s from different organisms have been used as adjuvants in vaccine formulations, such as fusion proteins, complexes peptides/HSP90, or as mixtures of peptides + HSP90[7]. Although the most used strategies in the design of vaccines based on HSP90 as adjuvants consist of the covalent union or formation of union complex between the antigenic peptides and these chaperones, the mix of the HSP90 with the antigens of interest is a valuable strategy[8–13]. Several works showed that antigenic peptides + HSP90 administration as a mixture are able to modulate the humoral and cellular immune responses produced against the antigens, improving protection against intracellular pathogens such as parasites and viruses[7,12,14–16]. Recently, results obtained by our group showed that different Hsp90 isoforms derived from plants differentially modulate immune response profiles. While the Hsp90 fusion strategy to the peptide would guarantee a Th1 immune response, the mixture of Hsp90 + peptide generates a Th1/Th2 immune response against the antigens[13,16,17]. Hsp90 from plants has the advantage of being a safe source with which humans have permanent contact. However, this advantage over other adjuvant systems still requires a greater understanding of the role and capabilities of plant Hsp90 (pHsp90) in immune response modulation. This result implies an increase in the number of vaccine models. Taking advantage of the fact that SARS-CoV2 is still a permanent infection on the world and a model of how the generation of alternative vaccines allowed us to control the effects of this pandemic, here, we propose the use of a short version of the Receptor Binding Domain (RBD) of the Spike protein of SARS-CoV2, the principal candidate in the design of subunit vaccines. We evaluated humoral and cellular immune responses against RBD through the strategy “protein mixture” (adjuvants + antigens). Furthermore, in this work, we analyzed the potential of the different formulations studied to neutralize viral infections.

## 2. Materials and Methods

### 2.1. Plasmid construction Receptor Binding Domain

The Receptor Binding Domain (RBD), residues 401 to 541, from Spike protein was reported of SARS-CoV2 Wuham isolate Wuhan-Hu-1, complete genome, NCBI, No. Accession 6XR8_A[18]. The *RBD* gene, was synthetized and codon optimized by GenScript Company (USA) in pUC57 plasmid. In addition, restriction sites 5’ *Pst*I and 3’ *Hind*III were added into the N-terminus and C-terminus, respectively. The recombinant plasmid was digested with *Pst*I and *HindI*II and the insert was directionally cloned into a 6xHis-tagged pRSET-A expression vector resulting in the pRSET-A-RBD plasmid.

### 2.2. Expression and purification of recombinant RBD

The pRSET-A-RBD plasmid was transformed in *E. coli* BL21 Star™(DE3) competent cells (Invitrogen). Bacteria were cultured to an OD_600_ of 0.5-0.6 and then protein expression was induced with isopropyl-β-d-thiogalactoside (IPTG) to a final concentration of 1 mM for 4-6 hours. Cells were harvested by centrifugation and stored at −20°C until use. Soluble RBD was purified under denaturing conditions using a nitrilotriacetic acid-Ni2+ column (Qiagen)[19]. Recombinant AtHsp81.2 and NbHsp90.3 were produced under native conditions as described in Corigliano et al.[19].

### 2.3. Mice and vaccination

C57BL/6 (H-2b) mice specific-pathogen-free were obtained from the central bioterium of Facultad de Ciencias Exactas y Naturales of Universidad de Buenos Aires. Six- to eight-week-old male and female mice were bred and housed following the institutional guidelines of the Universidad de General San Martín (C.I.C.U.A.E., IIB-UNSAM). Mice had access to food and water ad libitum and were maintained at 21-22°C, 12:12 h light-dark photocycle. Mice were randomly separated into seven groups and were vaccinated two times by intramuscularly (i.m.) injection at 21 days intervals (0 and 21 days post-vaccination, respectively) with equimolar doses of each recombinant protein as follows; Group PBS (negative control): 200 µl of sterile PBS, Group rNbHsp90.3: 6 µg of rNbHsp90.3 in 200 µl of sterile PBS; Group rAtHsp81.2: 6 µg of rAtHsp81.2 in 200 µl of sterile PBS, Group rRBD: 4 µg of rRBD in 200 µl of sterile PBS, Group rRBD + rAtHsp81.2: 4 µg of rRBD formulated with 6 µg of rAtHsp81.2 in 200 µl of sterile PBS, Group rRBD + rNbHsp90.3: 4 µg of rRBD formulated with 6 µg of rNbHsp90.3 in 200 µl of sterile PBS, and Group rRBD + Alum (positive control): 4 µg of rRBD formulated with 0.5 mg of Aluminum hydroxide in 200 µl of sterile PBS (Fig. 1d).

**Figure 1.**
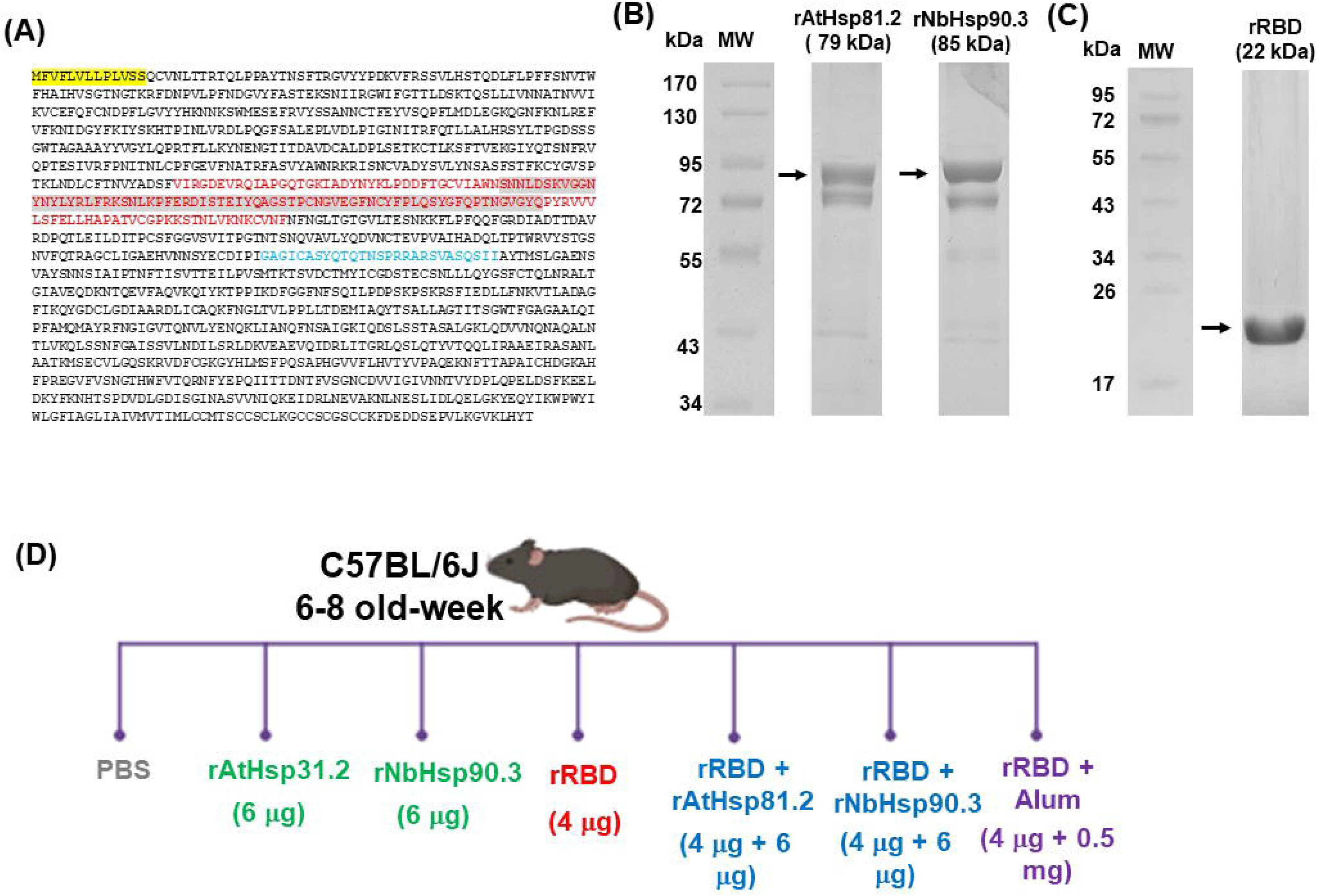
The solubilized and purified *Arabidopsis thaliana* Hsp81.2 (AtHsp81.2), *Nicotiana benthamiana* Hsp90.3 (NbHsp90.3), and RBD recombinant proteins. (A) The complete amino acid sequence of the Spike glycoprotein according to ACCESSION 6XR8_A (National Institutes of Health, National Center for Biotechnology Information database)^46^. The red letter indicates the region cloned to generate rRBD. Highlighted in gray is the ACE2 receptor binding RBD region^52^. Highlighted in yellow is the signal peptide. Highlighted in blue is the cleavage recognition sequence. (B) SDS-PAGE analysis under reducing conditions of rAtHsp81.2 and rNbHsp90.3 expressed and purified from *E. coli* Rosetta (DE3) pLys S. (C) SDS-PAGE analysis under reducing conditions of rRBD motif expressed and purified from *E. coli* BL21 pLys S. (D) C57BL/6 females and males mice were injected intramuscularly with four µg of rRBD alone (rRBD group) or formulated with six µg of rAtHsp81.2 (rRBD + rAtHsp81.2 group) or rNbHsp90.3 (rRBD + rNbHsp90.3 group) or with 0.5 mg of alum (rRBD + alum group) or PBS 1X (PBS group). MW: prestained molecular weight protein marker. Figure 1a was created with BioRender.com.

### 2.4. Antibody response and isotype determination

Anti-rRBD, anti-rAtHsp81.2 and anti-rNbHsp90.3 antibodies in the sera of vaccinated mice were determined by ELISA. Before each immunization, blood was collected from the cheek, and serum was stored at −20°C until analysis for the presence of specific antibodies. The sera were obtained at 0-, 21-, 42-, 63-, 84- and 105-days post-vaccination. Pre-immunization serum samples were taken for using as negative controls. Antigen-specific antibodies were analyzed by ELISA immunoassay as we described previously^48,49^. Flat-bottomed, polystyrene 96-well ELISA plates (ExtraGENE) were coated overnight at 4°C with 5 µg/ml of rRBD (IgGt, IgG1, IgG2b and IgG2a). To assess the immunogenicity of recombinant proteins for IgG production, mice serum antibodies were detected by indirect ELISA against specific recombinant proteins either anti-rRBD (1:500), followed by horseradish peroxidase (HRP)-conjugated anti-mouse IgG antibody (1:3000; Cell Signaling Technology Inc.). For isotype determination, mice sera were diluted 1:500 and rat anti-mouse IgG1, IgG2a or IgG2b horseradish peroxidase conjugated (1:10000; Sigma-Aldrich) were used as secondary antibodies. Tetramethyl-benzidine substrate (TMB; Invitrogen) was added and plates were read at 655 nm with an automatic ELISA reader (Synergy H1; Bio-Tek). All samples were measured by duplicate.

### 2.5. Cytokine analysis

The cytokine production was evaluated in spleens cells isolated from two or four immunized mice from each group 126 days post-vaccination. Splenocytes cultures (2 x 10^6^ cells/well) were stimulated with 10 µg/ml of rRBD and only medium as control non stimuli. Supernatants were harvested at 48 h to analyze IL-4 and IL-10 or 72 h for IFN-γ and stored at −70 ◦C until samples were measured by ELISA immunoassay as previously described in Corigliano et al.[20].

### 2.6. Co-Immunoprecipitation (Co-IP)

Co-IP assays were performed as described in Vanagas et al.[21] with minor modifications. Protease inhibitor cocktail (Sigma) was added in every step. An equimolar mixture of rRBD + rAtHsp81.2 or rRBD + rNbHsp90.3, or rRBD alone was incubated with protein A/G Plusagarose (sc-2003, Santa Cruz). Immunocomplexes were washed twice with washing buffer 1 (50 mM Tris, pH 8, 200 mM NaCl and 0.05 % Igepal100), twice with washing buffer 2 (50 mM Tris, pH 8, 300 mMNaCl and 0.05 % Igepal100), and twice with buffer TE (10 mM Tris, pH 8, 1 mM EDTA); then resuspended in 60 µl of SDS-PAGE loading buffer. Samples were boiled for 5 min and 20 µl were loaded per well in a 12 % SDS-PAGE gel for immunoblotting.

### 2.7. Pseudovirus neutralization assay

HEK293T cells (American Type Culture Collection (ATCC) CRL 3216) were cultured in DMEM with 10% FBS (Gibco) and 1% PenStrep (Gibco) with 5% CO2 in a 37 °C incubator (Thermo Fisher). A well-established pseudovirus neutralization system was adapted from[22,23]. In brief, pseudo-type lentiviruses coated with the SARS-CoV-2 S protein harboring the vector pCMV14-3X-Flag-SARS-CoV-2 S (a gift from Zhaohui Qian (Addgene plasmid #145780) were prepared by co-transfecting HEK293T cells with psPAX2 (a gift from Didier Trono (Addgene plasmid # 12260) and pLentipuro3/TO/V5-GW/EGFP-Firefly Luciferase (a gift from Ethan Abel (Addgene plasmid # 119816). Transfected HEK293T cells were incubated at 37 °C under 5% CO2 for 2 days, after which the supernatant containing pseudo-typed lentivirus particles was harvested and filtered through a 0.45-µm filter (Millipore). The viral particles were concentrated by centrifugation at 3000 x g overnight at 4°C and the pellet was resuspended in DMEM. hACE2-expressing HEK293 cells were seeded in 50 ml of medium in 96-well plates 1 day before transduction. Two-fold serial dilutions of sera were prepared in DMEN in a separate 96-well plate. The serum concentration ranged from 1/80 to 1/1280 diluted solution and incubated for 1 h at 37 °C with pseudotype Sars-CoV-2. After 48 h, the transduction efficiency was analyzed by quantifying luciferase activity. The 293T/17 cells for luciferase assay analysis were lysed by adding an equal amount of 2× lysis buffer (25 mM Tris hydrochloride pH 8, 2 mM DTT, 2 mM EDTA, 10% glycerol, and 1% Triton X-100) followed by incubation at room temperature on a shaker for 10 min. 20 μL of the cell lysates were added to a black 96-well FluoroNunc plate he luciferase reaction was initiated by the addition of 100 μL of the reaction buffer (25 mM Tricine hydrochloride pH 7.8, 5 mM magnesium sulfate, 0.5 mM EDTA, pH 8.0, 3.3 mM DTT, 0.5 mM ATP pH ∼7–8 (Sigma-Aldrich), 1 mg/mL BSA, 0.05 mg/mL D-luciferin (Gold Biotechnology), 0.05 mM co-enzyme A, and was read using a luminescence counter. Relative luciferase units were plotted and normalized in Prism (GraphPad) using a zero value of cells alone and a 100% value of 1:2 virus alone. Nonlinear regression of log(inhibitor) vs. normalized response was used to determine IC50 values from curve fits.

### 2.8. Statistical analysis

Statistical analysis was carried out with the Prism 5.0 Software (GraphPad) using two-way analysis of variance (ANOVA). Values of p < 0.05 were considered significant.

## 3. Results

### 3.1. The novel immune potent recombinant plant Hsp90 adjuvanted anti-COVID-19 vaccine, inducing RBD-specific antibodies in mice

Previously, Jangar et al.^18^ showed that the monomeric SARS-CoV2 spike protein receptor binding domain (RBD) has lower immunogenicity than the full-length spike (S) protein. Therefore, we selected RBD as an antigen for a better understanding of differentiated immune responses triggered by novel plant HSP90 (pHSP90) adjuvants. The RBD contains the region of the S protein necessary for binding to the human ACE2 receptor (hACE2) and for viral entry, and thus contains most epitopes targeted by neutralizing antibodies (nAbs) as well as multiple T cell epitopes[24–26].

The advantage of the recombinant protein strategy is that the region expressed can be chosen, minimizing undesirable regions. At the beginning of the project, it was still unknown whether undesirable epitopes of the spike protein could induce adverse immune responses. Therefore, we selected a minimal region containing the Receptor Binding Motif (RBM) that can efficiently generate neutralizing antibodies. In this way, we could reduce undesirable effects if they exist. Figure 1a shows the expressed RBD region (V401 to F541), which includes the ACE2 binding site (RBM) flanked by 32 and 30 residues at N- and C-terminal regions, respectively. The solubilized and purified *Arabidopsis thaliana* Hsp81.2 (AtHsp81.2), *Nicotiana benthamiana* Hsp90.3 (NbHsp90.3), and RBD recombinant proteins were observed in the purified fraction of SDS-PAGE (Fig. 1b and 1c). To evaluate whether rAtHsp81.2 and rNbHsp90.3 increase rRBD immunogenicity and modulate humoral response profile, we immunized C57BL/6J mice intramuscularly (i.m.) with 4 µg per mouse of rRBD formulated with 6 µg per mouse of rAtHsp81.2 or rNbHsp90.3 at Days 0 and 21 (Fig. 1d). Mice in the control groups received only rRBD (4 µg), only adjuvant (6 µg) or PBS. We immunized a seventh group with rRBD (6 µg) + alum (0.5 mg) as a positive control group.

After each injection and during 105 days at 21 interval-days, blood was collected and analyzed by an enzyme-linked immunosorbent assay (ELISA) using rRBD (Fig. 2a). Mice immunized with rRBD formulated with rAtHsp81.2 or rNbHsp90.3 adjuvants induced serum rRBD-specific IgG 42 days after the second immunization (Fig. 2b), with significantly higher IgG in groups that received adjuvanted versus unadjuvanted antigen. Notably, adjuvanted rRBD with alum induced lower antigen-specific IgG than adjuvanted antigen with pHsp90 (Fig. 2b). Interestingly, female and male mice immunized with rRBD + rpHsp90s did not show differences in the levels of anti-rRBD IgG. However, rRBD + alum-immunized female mice showed lower anti-rRBD IgG levels than male mice immunized with this formulation (Fig.2c and 2d). In addition, controls groups showed minimal antigen-specific IgG.

**Figure 2.**
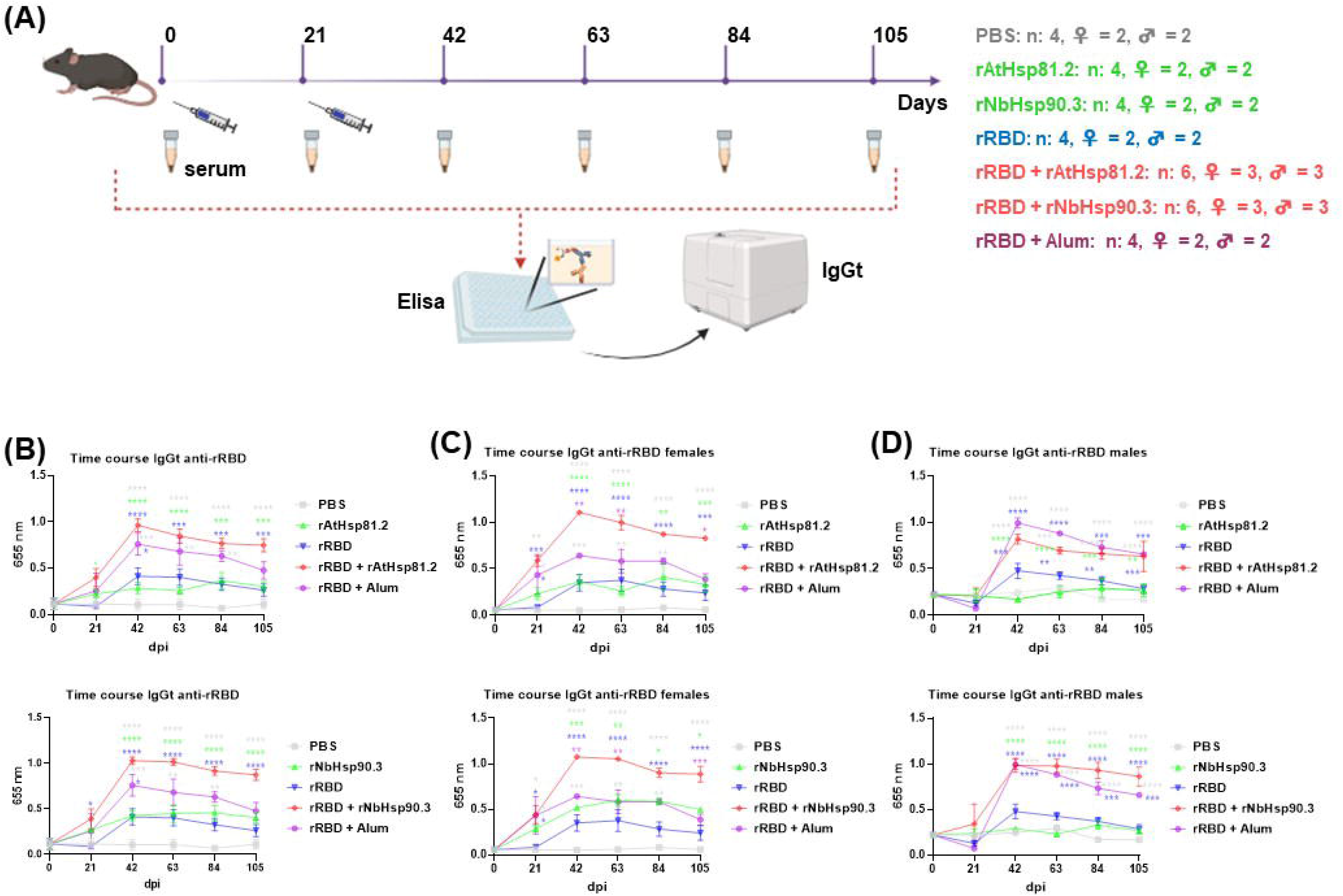
Two immunizations with rRBD adjuvanted with rAtHsp81.2 or rNbHsp90.3 are sufficient to induce a robust IgG total response in both female and male mice. (A) Study design including vaccination, sampling time points, and ELISA. The vaccination was performed twice at 21-day intervals via intramuscular route. Blood samples were collected at 0-, 21-, 42-, 63-, 84-, and 105-days post-vaccination in mice for serological assays. (B) rRBD-specific IgG levels in both female and male mice, in female mice (C) and male mice (D). Data are presented as mean ± SEM. Statistical analyses were performed using two-way ANOVA with Tukey’s multiple comparisons test. *p < 0.05; **p < 0.01; ***p < 0.001, and ****p < 0.0001 shown only for rAtHsp81.2/rNbHsp90.3/Alum + rRBD compared to other goups. Figure 2a was created with BioRender.com. A representative experiment of 3 independent replicates with similar results is shown.

To investigate the quality of the antibody responses induced, we measured IgG subtypes IgG2b, IgG2a, and IgG1 at day 84 as a representation of Th1 and Th2-type responses, respectively (Fig. 3a). Consistent with the literature, alum induces a higher IgG1 response compared to IgG2[27]. Even though we did not find a difference in the magnitude of the IgG responses between both rpHsp90s, strikingly, we did find a different profile of IgG2b/IgG2a versus IgG1 responses induced by these adjuvants (Fig. 3b, 3c, and 3d). Whereas rAtHsp81.2 induced significantly higher IgG1 responses compared to rNbHsp90.3 and non-adjuvanted RBD (Fig. 3b), rNbHsp90.3 induced significantly higher IgG2a/b responses, demonstrating a Th1-biased immune response (Fig. 3c and 3d). We further determined the ratio of IgG1/IgG2 responses, which showed that while alum induces a Th2-type response, AtHsp81.2 promotes a mixed Th1/Th2-type response, and NbHsp90.3 promotes a Th1-type response (Fig. 3b, 3c, and 3d).

**Figure 3.**
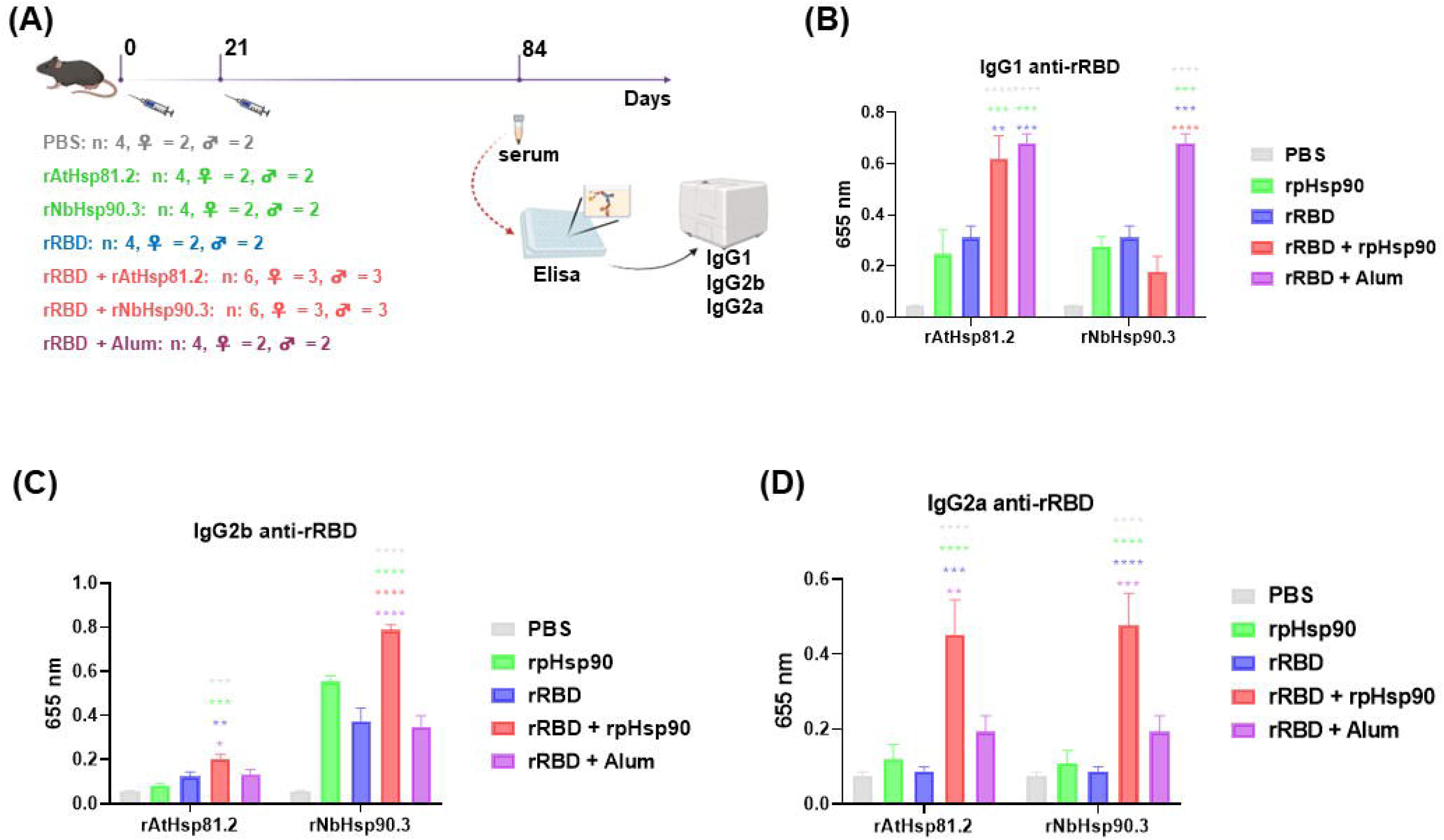
Different profile of humoral responses induced by rRBD adjuvanted with rAtHsp81.2 versus rNbHsp90.3. (A) Study design including vaccination, sampling time points, and ELISA. The vaccination was performed twice at 21-day intervals via intramuscular route. rRBD-specific IgG subclass antibodies for IgG1 (B), IgG2b (C), IgG2a (D), were measured at 84 days post-vaccination. Data are presented as mean ± SEM. Statistical analyses were performed using two-way ANOVA with Tukey’s multiple comparisons test. *p < 0.05; **p < 0.01; ***p < 0.001, and ****p < 0.0001 shown only for rAtHsp81.2/rNbHsp90.3/Alum + rRBD compared to other groups. Figure 3a was created with BioRender.com. A representative experiment of 3 independent replicates with similar results is shown.

### 3.2. RBD immunization with different recombinant plant Hsp90 adjuvants induces a distinctive profile of cytokines

To evaluate the cellular immune response induced by vaccination, we immunized two groups of four mice, each with rRBD adjuvanted with AtHsp81.2 or NbHsp90.3 with the same immunization regimen described above (Fig. 4a). Other two groups of two mice, each with non-adjuvanted rRBD and rRBD adjuvanted with alum, were included, as well as a PBS immunized control group (Fig. 4a). On day 126 post-first immunization, spleen cells were isolated and stimulated *in vitro* with rRBD, and IFN-γ, IL-10, and IL-4 production was detected by sandwich ELISA (Fig. 4b, 4c and 4d).

**Figure 4.**
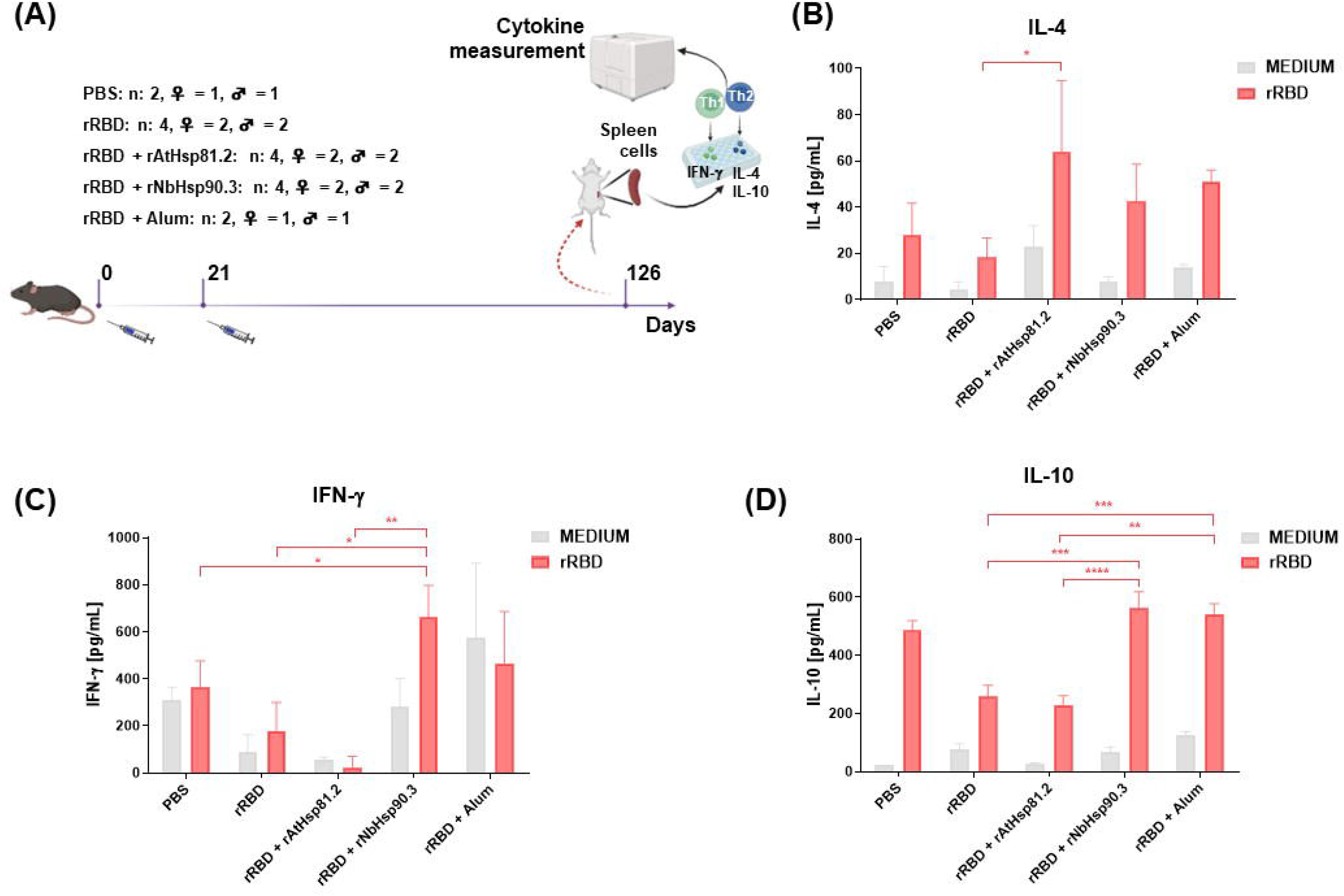
rRBD adjuvanted with rNbHsp90.3 or rAtHsp81.2 shows differences in cytokine profile. (A) Study design including vaccination, sampling time points, and cytokine measurement. At 126 days post-vaccination, levels of secreted cytokines were measured in the cell supernatant by sandwich ELISA for IL-4 (B), IFN-γ (C), IL-10 (D). Data are presented as mean ± SEM. Statistical analyses were performed using two-way ANOVA with Tukey’s multiple comparisons test. *p < 0.05; **p < 0.01; ***p < 0.001, and ****p < 0.0001 shown only for rAtHsp81.2/rNbHsp90.3/Alum + rRBD compared to other groups. Figure 4a was created with BioRender.com. A representative experiment of 2 independent replicates with similar results is shown.

We observed that splenocytes from mice vaccinated with rRBD adjuvanted with AtHsp81.2 secreted increased levels of IL-4 upon stimulation, a Th2 cytokine, compared to non-adjuvanted rRBD group (Fig. 4b). Interestingly, cells collected from the spleens of animals immunized with rRBD adjuvanted with NbHsp90.3 produced significantly higher, IFN-γ levels, a Th1 cytokine, compared to PBS, non-adjuvanted rRBD and rRBD adjuvanted with AtHsp81.2 groups (Fig. 4c). At the same time, NbHsp90.3 and alum adjuvanted groups showed a significant increase in IL-10 secretion compared to non-adjuvanted rRBD and rRBD adjuvanted with AtHsp81.2 groups (Fig. 4d). In addition, stimulated splenocyte from mice from all the included groups induced significant IL-10 secretion compared to non-stimulated splenocytes (Fig. 4d). In summary, *in vivo* studies in the mouse model demonstrated that adjuvating rRBD (in low dose) with rNbHsp90.3 or rAtHsp81.2 induced potent humoral and cellular immune responses, but with differences in the cytokine profile elicited, that were higher than those elicited by the formulation adjuvanted with alum.

### 3.3. Vaccination with rRBD adjuvanted with NbHsp90.3 but not rRBD adjuvanted with AtHsp81.2 elicits SARS-CoV2 neutralizing antibody responses

To investigate the potential of each pHsp90 adjuvant to enhance protection against infection with a SARS-CoV2 variant, we evaluated the virus neutralization capability of antibodies induced by the rRBD adjuvanted with AtHsp81.2 and NbHsp90.3. Neutralizing antibody (nAb) titers were measured in the serum from immunized mice 42 days after the first immunization using a lentivirus-based pseudovirus (PSV) neutralization assay (Fig. 5a). Sera were incubated with entry into hACE2 expressing HEK293T cells and quantified as a function of the luciferase reporter gene transduction. We found that while both adjuvants enhanced the immunogenicity of rRBD, each adjuvant formulation showed differences in neutralizing antibody capacity. and less so in the rRBD + rAtHsp81.2 group (Fig. 5b and 5c). In the alum group, we did not detect neutralizing-antibody responses. The rRBD adjuvanted with rNbHsp90.3 group had high nAb titers (1:365) and also significantly higher titers than the alum, non-adjuvanted antigen, and PBS groups (Fig. 5b). In addition, nAbs (<1:160) against the PSV were detected in at least some animals in rRBD adjuvanted with AtHsp81.2 group (Fig. 5c). While rRBD + rNbHsp90.3 and rRBD + rAtHsp81.2 formulations elicited similar rRBD-specific IgG levels, the higher nAb titers observed with rNbHsp90.3 suggest improved antibody quality.

**Figure 5.**
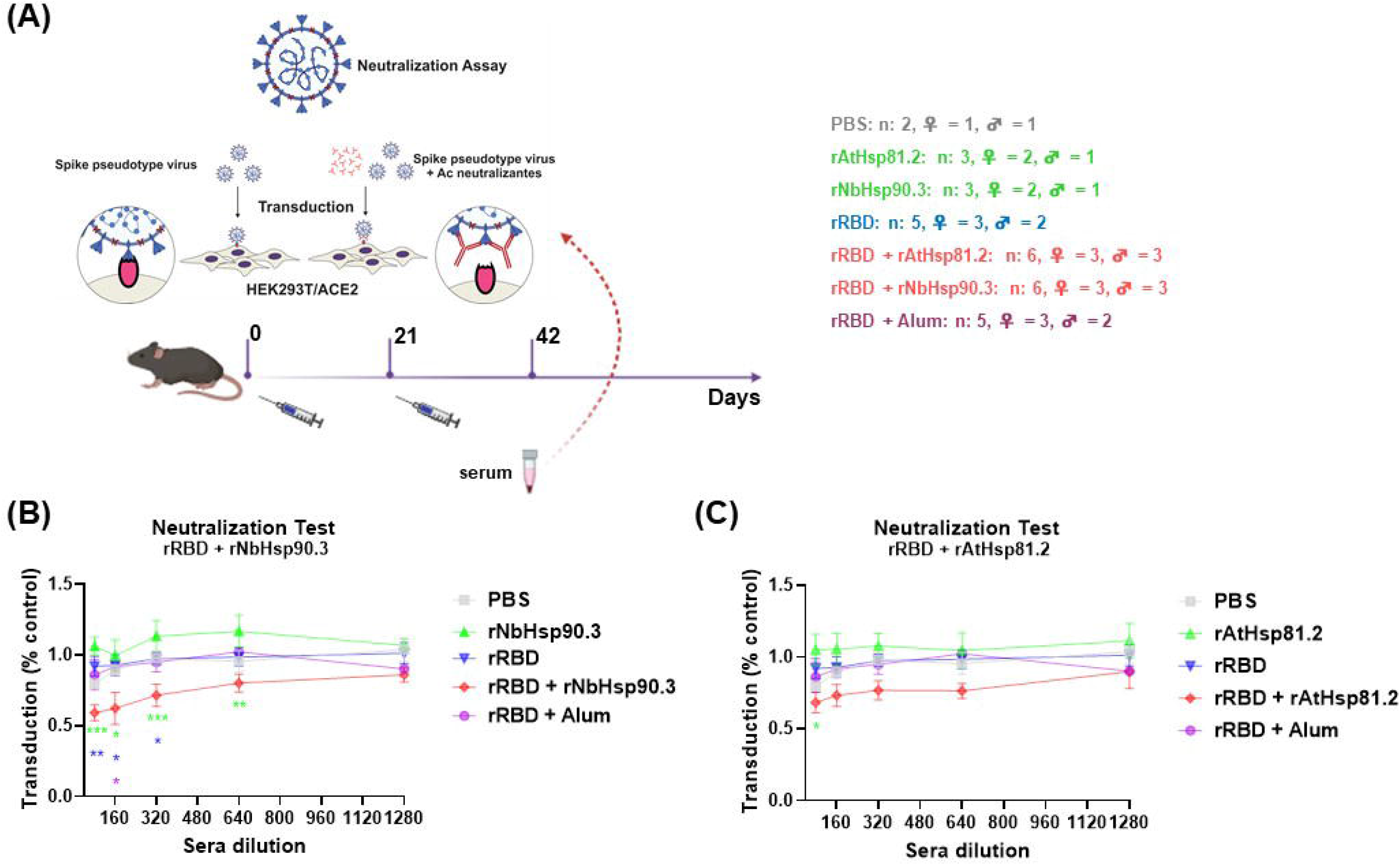
rRBD adjuvanted with recombinant plant Hsp90 elicites a neutralizing antibody response. (A) Schematic diagram of the vaccination and neutralization study. Anti-RBD neutralizing activity by virus neutralization tests in rRBD + rNbHsp90.3 group (B), and rRBD + rAtHsp81.2 group (C) 42 days post-immunization. Data are presented as the mean ± SEM. Statistical analyses were performed using a two-way ANOVA with Tukey’s multiple comparison test. *p < 0.05; **p < 0.01; and ***p < 0.001 shown only for rAtHsp81.2/rNbHsp90.3 + rRBD compared to other groups. Figure 5a was created using BioRender.com. A representative experiment with independent replicates and similar results is shown.

### 3.4. Complex assembly of rRBD with rNbHsp90.3, but not with r AtHsp81.2

An intriguing question of our model is whether both chaperones must form complexes to generate an effective immune response that affects the accompanying antigens[7]. With this idea in mind, we performed co-immunoprecipitation (co-IP) assays. rNbHsp90.3 or rAtHsp81.2 were mixed separately with the rRBD antigen and, after incubation, were co-IP with their respective antibodies (Fig. 6a). Figure 6b shows that rRBD interacted with rNbHsp90.3, but not with rAtHsp81.2 confirming the assembly of the rRBD/rNbHsp90.3 complex. Co-IP performed by using rRBD alone as a control shows no reactivity with any of the antibodies assayed (Fig. 6c). These data indicate that the greater neutralizing capacity observed in the rRBD adjuvanted with rNbHsp90.3 group would be given by the rRBD-rNbHsp90.3 interaction rather than by the quality of the immune response triggered by the adjuvants.

**Figure 6.**
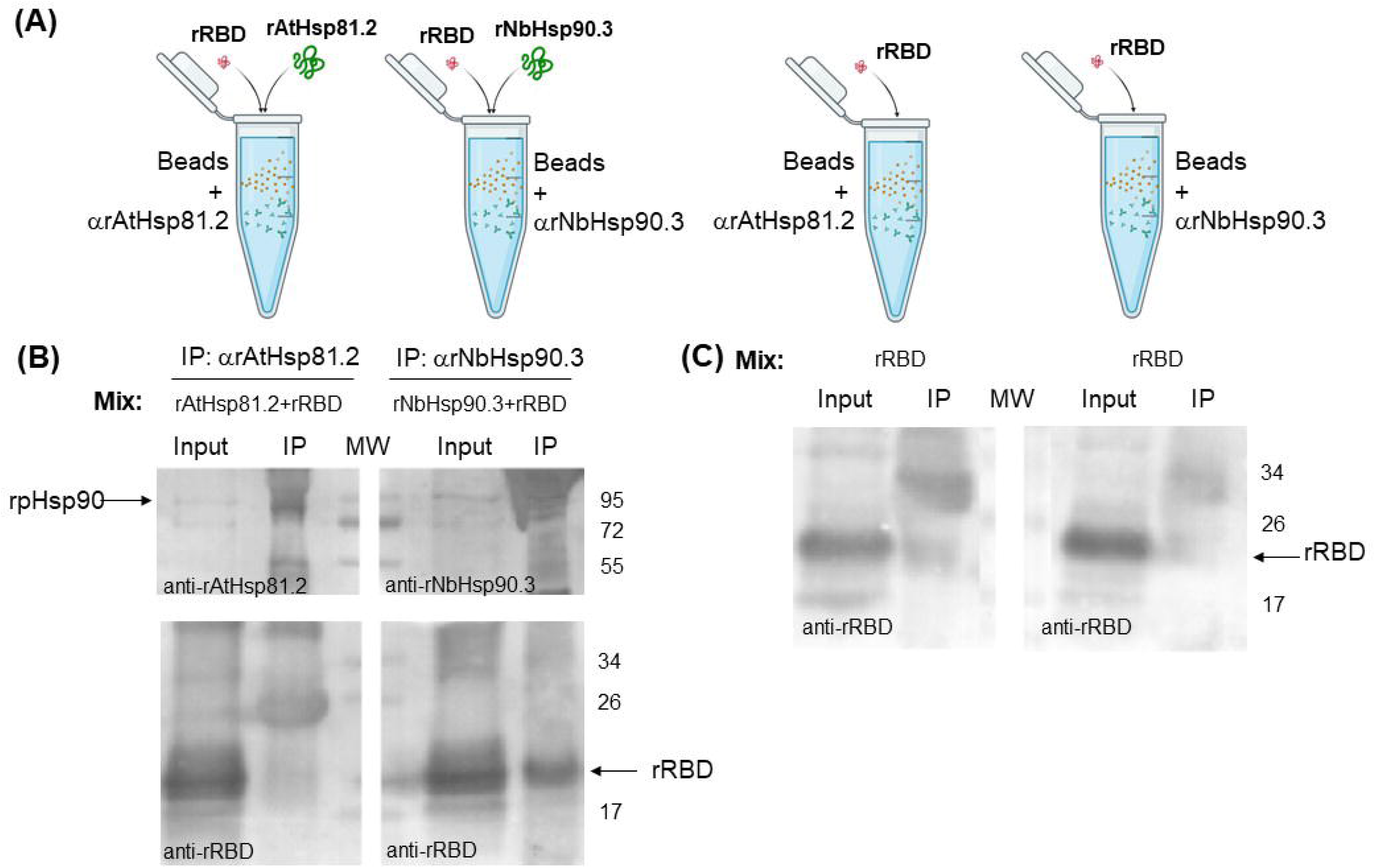
rRBD interacted with rNbHsp90.3, but not with rAtHsp81.2. (A) Schematic diagram of co-immunoprecipitation (co-IP) assays. Co-immunoprecipitation-WB analysis from rRBD + rNbHsp90.3, rRBD + rAtHsp81.2 (B) or rRBD alone (C). A representative Western-blot is shown for anti-rNbHsp90.3, anti-rAtHsp81.2 and anti-rRBD. IP: immunoprecipitation. MW: prestained molecular weight protein marker. Figure 6a was created with BioRender.com. A representative experiment of 3 independent replicates with similar results is shown.

## 4. Discussion

The pandemic produced by SARS-CoV2 showed the importance of having a massive battery of safe immunization systems to respond quickly and stop its spread, saving a large proportion of the population from death, alleviating the health-system collapse, and breaking off the economic debacle. It is worth mentioning that in a record time, a large set of vaccines were approved (less than a year) throughout 2020, and around the world, nearly 11 billion vaccines were administered.

Although previous studies carried out on vaccines against Ebola or Mers-CoV have contributed to laying the foundations for vaccine development in pandemic situations[28], the pandemic experience with SARS-CoV2 consolidated the concept of rapid approval of vaccines through the intervention of international organizations such as the Food and Drug Administration (FDA) and the European Medicines Agency (EMA) for health emergencies[29]. In fact, a vaccine against SARS-CoV2 named ArVac-CG, based on a recombinant RBD antigen, was recently approved in Argentina[30]. Although this establishes solid foundations for future pandemic experiences that could take place in Argentina, emerging and socially and economically vulnerable countries remain at a disadvantage if new pandemic situations happen again. For this reason, the production of vaccines and their equitable distribution worldwide will require more significant deployment than those observed during this pandemic. In this sense, we consider it essential to continue incorporating safe vaccine models that can be quickly approved and that can also be produced in large quantities at acceptable costs.

Hsp90s have mainly contributed to improving vaccine development against infectious diseases, especially against intracellular pathogens. Several reports have demonstrated that Hsp90s from different sources are potent adjuvants, generating an appropriate immune response against infectious diseases[7]. Here, we showed that plant Hsp90 can be used as an efficient adjuvant to stimulate an effective immune response against a recombinant form of the SARS-COV2 RBD antigen. We observed that NbHsp90.3 generated the production of neutralizing antibodies and a Th1-type cellular immune response with the production of IFN-γ. On the contrary, AtHp81.2 induced a mixed Th2 humoral immune response with IL-4 production. In general, Hsp90s derived from different organisms, including those from plants, had already been shown to have immunogenic capacities and immunomodulatory properties^7^ but had not yet been studied as adjuvants for SARS-COV2. The fact that both pHsp90s generated different types of immune responses could be an advantage when choosing one or the other adjuvant. One of the most commonly used adjuvants for vaccine development in emergencies, which is allowed for use in humans, is aluminum salts, which are also widely used in vaccination for COVID-19[3]. Aluminum salts act stimulating immune responses, especially of the Th2 type[31,32], related to the B lymphocyte stimulation for antibody production independent of TLR and CD4+ T helper cell responses[33]. In general, this kind of activation is poor[34]. Therefore, alum adjuvants are used with other salts or adjuvants to enhance the response. For emergencies, a combination of imidazoquinoline class molecules (TLR7 and TLR8 agonist) adsorbed onto alum has also been approved for use in humans[35]. This combination facilitates the generation of cell-mediated immunity[36,37]. In addition, other adjuvants used with recombinant proteins for anti-COVID-19 vaccines are CpG, SQBA, AS03, and MF59, among others[3,22]. The advantage of using a plant version of Hsp90 as an adjuvant is that this protein is a natural plant compound with which humans can maintain permanent contact without showing toxicity. pHsp90s have been shown to stimulate the humoral and cellular immune response through cross-presentation by the internalization of exogenous Hsp90s complexed to or fused to a peptide in early endosomes and the induction of inflammatory cytokines via TLR4 and TLR2[7]. Similarly, pHsp90 has been shown to stimulate humoral and cellular responses in other models[13,16] in addition to interacting with TLR4 to mediate MHC I activation[7,20]. This would provide a new type of adjuvant to expand the vaccine production possibilities in general and in future pandemics.

As far as we know, the RBD protein (V401; F541) analyzed in this work is a shorter version than others previously studied. Mainly, it is smaller in the N-terminal region than several of those already used, such as NARUVAX-C19 (Q321, S521)[38], ArVac-CG (319R, 537K)[30] or (R328, T531)[22]. All of these were effective in eliciting a protective immune response. Once again, it shows the versatility of recombinant techniques to design responsive antigens limited to the region of interest. Our strategy was to present the RBD to the immune system to obtain neutralizing antibodies. However, we also detected a cellular response in mice immunized with rRBD adjuvanted with NbHsp90.3, even though no T epitopes have been identified in this fragment during natural infection[39]. This fact may suggest that Hsp90 could help antigen presentation exogenously[40,41] or through the internalization of the peptides from the endosome to the cytosol by the proteasome to the re-presentation of Hsp90-associated peptides[7,42], at the same time that Hsp90s *per se* could trigger the secretion of cytokines.

Hsp90s are specialized chaperones that can bind to a group of client proteins that are not necessarily unfolded[43]. Therefore, it is unknown whether its role as an adjuvant is due to its intrinsic capacity to stimulate the immune response in the formulation or is due to its ability to bind the immunogen as occurs in cancer models[44–48]. Here, we observed that of the two chaperones, only NbHsp90.3 would form a complex with rRBD, while the AtHsp81.2 + rRBD formulation would only be a mixture. Interestingly, the rRBD/NbHsp90.3 complex induced an immune response towards a Th1 profile, which correlates with the immunomodulatory properties commonly described for Hsp90s[7], while the formulation with AtHsp81.2 would elicit an immune response towards a Th2 profile, which is likely for a recombinant antigen alone. This implies that although this mixing strategy has the advantage of being quickly developed, it will be necessary to guarantee that the antigen and the adjuvant form complex in future vaccine formulations if a Th1 profile is required. However, we cannot rule out that these differences in the profile of the immune response observed are related to differences in the intrinsic properties of each chaperone or to the structure and characteristics of each antigen. Likewise, in cases where the antigen and the adjuvant do not generate a complex or the antigen/adjuvant mixture does not trigger a potent protective response, another alternative is the “fusion protein” strategy. However, it is more laborious, and the expression levels of the recombinant proteins vary from case to case. In this case, it was also shown to be highly efficient in generating an adequate immune response[7,13,17].

## 5. Conclusion

The emergence of the SARS-CoV-2 pandemic was a major global challenge for the entire health, research, and vaccine production systems in record time. Initially, numerous studies briefly addressed the lack of knowledge about the characteristics of the infection. Soon, it was clear that the vaccine would be a fundamental tool to control the infection. Fast track approval has promoted the use of different vaccines. In that sense, there is some uncertainty about the efficiency and feasibility of a safe vaccine. Fortunately, all the vaccines generated and approved have shown to be protective and with few adverse effects. However, having several vaccine systems is advantageous to enable a better response for this type of situation. In addition to the immunogenic properties of pHsp90s being similar to those observed in other Hsp90s, they would be a safe system for humans since it is a chaperone with which, as already mentioned, there is permanent contact through food. We were able to show that pHsp90 can indeed be an adjuvant to take into account for the development of anti-COVID-19 vaccines. Interestingly, we also show that these properties may be related to their ability or not to form complexes with the antigen of interest, which should be analyzed on a case-by-case basis.

## Author contributions

M.C. and S.O.A. designed and coordinated the research and analyzed data. V.A.R.D. performed all experiments and analyzed data. A.A., L.F.M.M and M.A.S. helped to perform the recombinant protein purifications. C.J.G. and A.O. performed the neutralizing-antibody response assays and helped to interpret experiments and analyze data. M.G.C. and V.A.S. helped to perform the cytokine analysis. A.L. performed the mice immunizations. All authors wrote the initial draft, and all authors contributed to the final manuscript.

## Institutional Review Board Statement

All mice were maintained under specific-pathogen-free conditions and handled according to the approved institutional animal care and use committee protocols of the Universidad de General San Martín (C.I.C.U.A.E., IIB-UNSAM, 09/2016).

## Informed Consent Statement

Not applicable.

## Data Availability Statement

All data upon which conclusions are drawn are included in the manuscript.

## Acknowledgements

S.O.A. Angel, M. Clemente, M.G. Corigliano and V.A. Sander are members of Consejo Nacional de Investigaciones Científicas y Técnicas (CONICET) and Professor of Universidad Nacional General San Martin (UNSAM). C. Jaquenod de Giusti and A. Orlowski are members of CONICET and Professor of Universidad Nacional de La Plata (UNLP). V.A. Ramos-Duarte and L.F. Mendoza-Morales are fellows of CONICET. M.A. Sáchez is fellow of Agencia Nacional de Promoción de Ciencia y Técnica (ANPCyT). A. Legarralde is technical staff of CONICET. A. Atela is student of UNSAM.

